# Structure-function relationship for a divergent Atg8 protein required for a non-autophagic function in malaria parasites

**DOI:** 10.1101/2021.05.24.445495

**Authors:** Marta Walczak, Thomas R. Meister, Yili Zhu, Ellen Yeh

## Abstract

Atg8 family proteins are highly-conserved eukaryotic proteins with diverse autophagy and non-autophagic functions in eukaryotes. While the structural features required for conserved autophagy functions of Atg8 are well-established, little is known about the molecular changes that facilitated acquisition of divergent, non-autophagic functions of Atg8. The malaria parasite *Plasmodium falciparum* offers a unique opportunity to study non-autophagic functions of Atg8 family proteins because it encodes a single Atg8 homolog whose only essential function is in the inheritance of an unusual secondary plastid called the apicoplast. Here we used functional complementation to investigate the structure-function relationship for this divergent Atg8 protein. We showed that the LC3-interacting region (LIR) docking site (LDS), the major interaction interface of Atg8 protein family, is not sufficient for *Pf*Atg8 apicoplast function. Other regions previously implicated in canonical Atg8 interactions, the ubiquitin-interacting motif (UIM) docking site (UDS) and the *N*-terminal helix are not required for *Pf*Atg8 function. Finally, the unique Apicomplexan-specific loop previously implicated in interaction with membrane conjugation machinery *in vitro,* is not required *in vivo* neither for membrane conjugation nor for the effector function of *Pf*Atg8. These results suggest that the effector function of *Pf*Atg8 is mediated by structural features distinct from those previously identified for macroautophagy and selective autophagy functions.

**Importance:** The most extensively studied role of Atg8 proteins is in autophagy. However, it is clear that they have other non-autophagic functions critical to cell function and disease pathogenesis yet understudied compared to their canonical role in autophagy. Mammalian cells contain multiple Atg8 paralogs that have diverse, specialized functions. Gaining molecular insight into their non-autophagic functions is difficult because of redundancy between the homologs and their role in both autophagy and non-autophagic pathways. Malaria parasites such as *Plasmodium falciparum* are a unique system to study a novel, non-autophagic function of Atg8 separate from its role in autophagy: They have only one Atg8 protein whose only essential function is in the inheritance of the apicoplast, a unique secondary plastid organelle. Insights into the molecular basis of *Pf*Atg8’s function in apicoplast biogenesis will have important implications for the evolution of diverse non-autophagic functions of the Atg8 protein family.

## Introduction

Atg8 family proteins are highly conserved eukaryotic proteins with important functions in autophagy and non-autophagic pathways in eukaryotes (1, 2). This small, ubiquitin-like protein is covalently conjugated to membrane lipids where it recruits effectors, lying at the nexus of protein interactions to coordinate membrane dynamics (1, 3, 4). Although its role in autophagy is the best defined, Atg8 family proteins are functionally diverse. Atg8 and its conjugation machinery (Atg4, Atg3, and Atg7) are ubiquitous among eukaryotes, suggesting that autophagy or another Atg8-associated membrane function was a core pathway present in the last common eukaryotic ancestor (5, 6). From this highly conserved starting point, the single ortholog found in fungi and protists has expanded to multiple Atg8 paralogs in mammals and plants, hypothesized to represent functional specialization and diversification (7).

In humans at least six Atg8 homologs have been identified: LC3A, LC3B, LC3C, GABARAP, GABARAPL1/GEC1, and GABARAPL2/GATE-16 play partly redundant roles in autophagy but also participate in a number of non-autophagic pathways such as phagocytosis, vesicle trafficking, secretion, and exocytosis that do not overlap across homologs (2, 3, 8). These non-autophagic functions highlight the integral role of the Atg8 protein family in the mammalian endomembrane system beyond their role in autophagy. Unfortunately, overlapping autophagy and non-autophagic functions of mammalian Atg8 homologs make it challenging to investigate the functional diversification of this conserved protein family.

Atg8 proteins have also evolved a non-autophagic function in a distinct endomembrane compartment in a group of divergent eukaryotes. *Plasmodium* spp, the protozoa that cause malaria, and related apicomplexan parasites such as *Toxoplasma gondii* contain a non-photosynthetic plastid derived from secondary endosymbiosis called the apicoplast. During secondary endosymbiosis, a chloroplast-containing alga was phagocytosed by another eukaryote and established as a multi-membraned plastid that is part of the endomembrane system (9, 10). Unexpectedly, the single Atg8 homolog in *T. gondii* was shown to be conjugated to the outer membrane of the apicoplast (*11*). In *Plasmodium*, it is unclear if macroautophagy occurs since core components of the initiation steps and typical lysosomes are missing (12). Instead, knockdown of Atg8 in *P. falciparum* causes loss of the apicoplast during cell division and parasite death. When apicoplast function is rescued in Atg8-deficient parasites, parasite growth is recovered, demonstrating that *Pf*Atg8’s novel function in apicoplast inheritance is its only essential function (13).

*Pf*Atg8’s novel apicoplast function offers a unique opportunity to investigate Atg8 protein structure-function relationships for a non-autophagic function in the absence of autophagy. Atg8 homologs are compact proteins of <150 amino acids. Previous protein structure-function studies have defined several Atg8 protein regions important for autophagy: 1) an Atg8-interacting motif/ LC3-interacting region docking site (LDS) that is a key binding pocket for cargo receptors and core autophagy machinery, 2) an *N*-terminal helical extension, a hallmark of Atg8 proteins distinguishing them from other members of ubiquitin superfamily, important for oligomerization, membrane tethering, and fusion, and most recently 3) a ubiquitin-interacting motif docking site (UDS) which interacts with several new autophagy receptors (14–17). These structure-function relationships have provided important molecular insight into functional specialization for selective autophagy, for example mutation of the LDS to modulate affinity and selectivity for specific cargo receptors (18, 19).

A significant gap in our knowledge is the molecular determinants important for non-autophagic functions. Do effectors also engage known LDS or UDS sites for non-autophagic functions? Is membrane tethering and fusion by the *N*-terminal extension selective for certain endomembrane compartments? More broadly, how have non-autophagic functions contributed to functional diversification and expansion of Atg8 protein families? Deciphering the molecular basis of *Pf*Atg8’s non-autophagic function in the apicoplast will reveal molecular evolution that facilitated functional diversification in this important protein family.

## Results

### Atg8 homologs with specialized autophagic and non-autophagic functions cannot complement the apicoplast function of *Pf*Atg8

Atg8 homologs from distantly-related species often can functionally complement autophagic functions in yeast and/or mammalian cells. *Pf*Atg8 as well as Atg8 homologs from *Leishmania major*, *Trypanosoma cruzi* and *Arabidopsis thaliana* have been able to at least partly restore autophagic function in *atg8Δ* yeast (20–23). These functional complementation experiments suggest that the structural features required for autophagic function are highly conserved among Atg8 homologs.

*Pf*Atg8 shares 33-43% sequence identity and 50-80% similarity with yeast and mammalian Atg8 homologs (Fig. S1). While the function of yeast Atg8 is limited to autophagy pathways, mammalian homologs show functional diversification. This is likely due to structural differences which affect the selectivity and specificity of their interactions in both autophagic and non-autophagic function. We wondered whether mammalian Atg8 homologs that showed functional diversification in autophagy and non-autophagic pathways could complement *Pf*Atg8’s function in the apicoplast. Episomes expressing Atg8 homologs from these different species were introduced into a *Pf*Atg8-TetR/DOZI strain, in which *Pf*Atg8 expression is regulated by anhydrotetracycline (ATc) via the TetR/DOZI repressor (13, 24). Each transgenic Atg8 homolog was expressed as an *N*-terminal GFP fusion protein and was truncated to remove the final amino acids in the native sequence exposing the terminal Gly residue that is conjugated to lipids (4). These constructs mimic the membrane conjugation of *Pf*Atg8, whose native sequence ends in a terminal Gly residue and does not require processing by *Pf*Atg4 prior to membrane conjugation (13, 25). Successful transfectants were obtained for episomes expressing: *Pf*Atg8WT (positive control), *Pf*Atg8 G124A mutant (negative control which cannot be conjugated to the membrane), *Sc*Atg8, and human LC3A, LC3B, LC3C and GABARAPL2/GATE-16. While transgenic GFP-*Pf*Atg8WT was able to fully restore parasites’ growth upon loss of *Pf*Atg8 expression, neither yeast Atg8 nor human LC3B, LC3C or GABARAP-L2 were able to restore growth in the absence of *Pf*Atg8 expression (Fig 1A and 1B). Parasites expressing GFP-LC3A showed a weak rescue phenotype with 28% growth compared to control parasites (grown in the presence of endogenous Atg8) after 4 reinvasion cycles.

**Figure 1.**
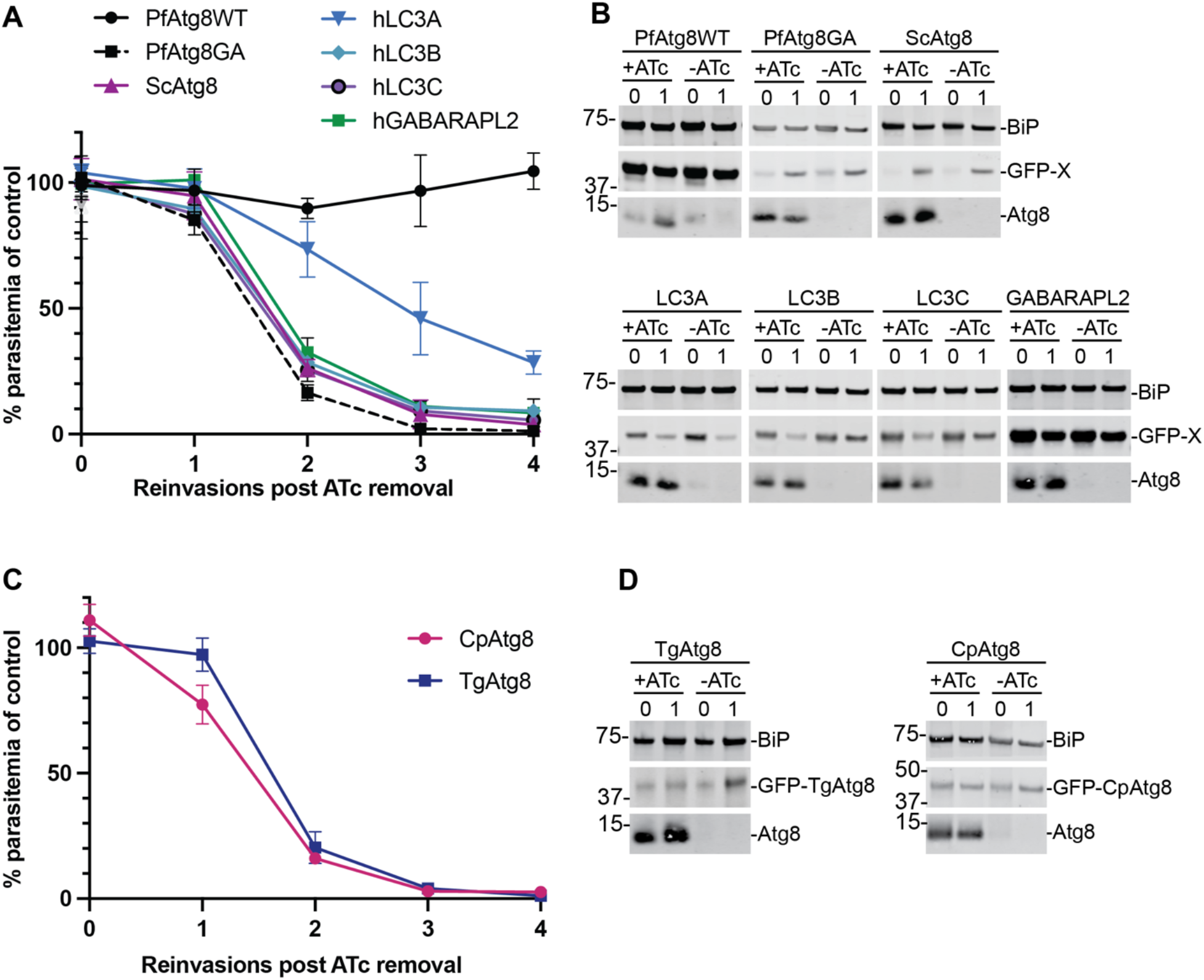
Atg8 homologs from other species do not efficiently complement *Pf*Atg8 function. (A) Growth of Atg8-TetR/DOZI parasites expressing indicated GFP-tagged homologs from human and yeast *S. cerevisiae* (h and Sc, respectively) in the absence of ATc over 4 reinvasion cycles. Parasitemia for each time point was normalized to the culture grown in the presence of ATc. Mean ± standard deviation of minimum 2 biological replicates is shown. (B) Western blots showing expression of indicated proteins during the first two cycles of the growth time course. Equal parasite numbers were loaded per lane. BiP, loading control; X, Atg8 homolog. Numbers above the blots represent parasite reinvasion cycles post ATc removal. (C) Growth of Atg8-TetR/DOZI parasites expressing *Tg*Atg8 or *Cp*Atg8 in the absence of ATc over 4 reinvasion cycles. Parasitemia for each time point was normalized to the culture grown in the presence of ATc. Mean ± standard deviation for 2 biological replicates is shown. (D) Western blots showing Atg8 knockdown and expression of GFP-*Tg*Atg8 or GFP-*Cp*Atg8 during the first two cycles of the growth time course. Equal parasite numbers were loaded per lane. BiP, loading control. Numbers above the blots represent parasite reinvasion cycles post ATc removal.

*Toxoplasma gondii* and *Cryptosporidium parvum* are apicomplexan parasites related to *Plasmodium* spp. *T. gondii* also has an apicoplast and *Tg*Atg8 is required for its inheritance, suggesting a similar non-autophagic function for *Pf*Atg8 and *Tg*Atg8 (11). *C. parvum*, on the other hand, lost the apicoplast in the course of evolution but retained a copy of the Atg8 gene (9). Both *Tg*Atg8 and *Cp*Atg8 have >60% sequence identity to *Pf*Atg8 (Fig. S1). We were interested if the Atg8 homologs from either of these two closely-related parasites can functionally complement *Plasmodium* Atg8. To this end, episomes containing GFP-*Tg*Atg8 or GFP-*Cp*Atg8 terminated with Gly residue were transfected into the *Pf*Atg8-TetR/DOZI strain. Surprisingly, neither of the homologs could restore growth in *Plasmodium* lacking endogenous Atg8 (Fig. 1C and D). Furthermore, only one out of three transfections with GFP-*Tg*Atg8 was successful, suggesting that the construct may be toxic to *Plasmodium*.

Altogether, these data suggest that *Plasmodium* Atg8 has structural features absent from other, even closely-related homologs, that are specifically required for its function in the *Plasmodium* apicoplast. The lack of efficient complementation of *Pf*Atg8 function by yeast, mammalian and apicomplexan homologs could be due to protein expression, misfolding or degradation, loss of membrane conjugation or, once attached to the membrane, inability to interact with effectors.

### Atg8 conjugation to the apicoplast membrane is specific but does not depend on the Apicomplexan-specific loop

The apicoplast function of *Pf*Atg8 requires its recognition and attachment to the apicoplast membrane via the action of membrane conjugation machinery, Atg3, Atg7, and a noncovalent Atg12-Atg5 complex (26, 27). Since none of the Atg8 homologs we tested could fully complement *Pf*Atg8, we screened these homologs for membrane association and apicoplast localization. To determine membrane association, cells expressing GFP-Atg8 fusions were subjected to Triton X-114 fractionation. This assay utilizes temperature-dependent phase separation of Triton X-114 into a detergent-rich fraction containing membrane-bound proteins and an aqueous fraction containing soluble and peripheral membrane proteins which are then analyzed by western blot (28). Because partitioning of proteins between the two fractions was variable between experiments, we normalized the percentage of membrane-bound GFP-Atg8 homolog fusions to the percentage of membrane-bound endogenous Atg8. To determine apicoplast localization, GFP-tagged Atg8 homologs were analyzed by live microscopy. Of all tested homologs, only *Cp*Atg8 showed membrane association and correct apicoplast localization, similar to *Pf*Atg8 (Fig. 2A-B).

**Figure 2.**
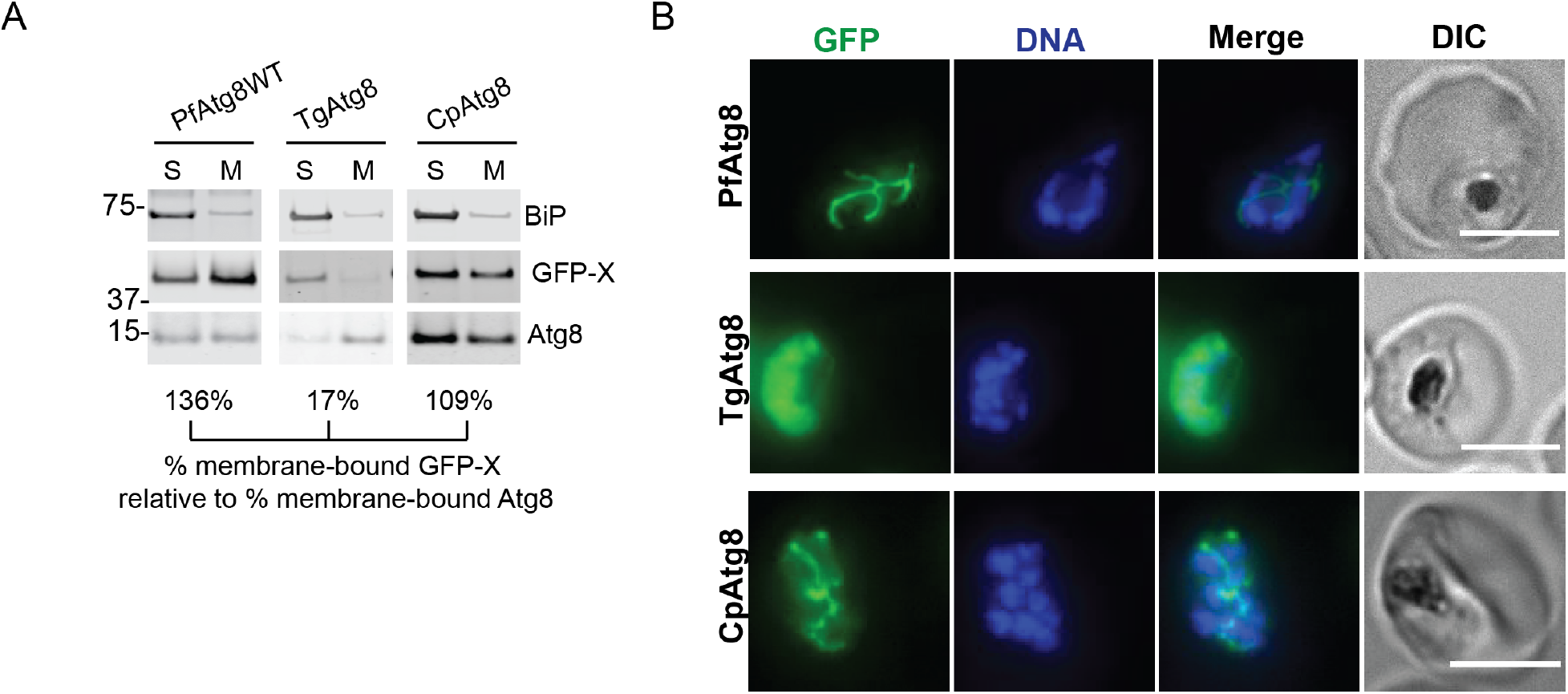
*Cp*Atg8, which does not functionally complement *Pf*Atg8 function, correctly localizes to the apicoplast membrane. (A) Western blot showing membrane association of selected proteins in a Triton X-114 fractionation experiment. Numbers below the blots represent the percentage of membrane-bound GFP-tagged Atg8 homolog normalized to percentage of the membrane-bound endogenous Atg8. S, soluble fraction; M, membrane fraction; BiP, soluble protein; Atg8, membrane-bound protein; X, Atg8 homolog. (B) Representative live fluorescence images showing the localization of selected GFP-tagged *Pf*Atg8 homologs. DNA was stained with Hoechst 33342. Maximum intensity projections are shown. Scalebar 5 μm.

The high specificity of Atg8 conjugation to the apicoplast membrane indicates that distinct structural features may govern recognition by the *Plasmodium* Atg8 conjugation machinery. Previous *in vitro* binding studies showed that interaction with *Pf*Atg3 was dependent on an extended loop region in *Pf*Atg8 (29). In the *Pf*Atg8 crystal structure, this 9 amino acid insertion into the loop between helix α3 and strand β5 within the canonical ubiquitin fold forms a hairpin that has not previously been seen in any other homologous Atg8 structures (29) (Fig. 3A-B). Notably, Atg8 homologs and putative homologs from other apicomplexan parasites and chromerids which possess a secondary plastid also contain an insertion in this loop region, designated the Apicomplexan-specific loop, though the sequence is not conserved (Fig. S2). To determine whether the extended loop motif is necessary for membrane conjugation of *Pf*Atg8, we transfected *Pf*Atg8-TetR/DOZI strain with an episome expressing *N*-terminal GFP-fusion of *Pf*Atg8 in which the 9 additional amino acids (amino acids 69-77) were removed (*Pf*Atg8ΔLoop). In addition, since Atg8 proteins share high structural homology and because deletion constructs may be disruptive to protein folding, hybrids of *Pf*Atg8 and a canonical homolog *Sc*Atg8 exchanging their respective loop regions, amino acids 69-82 in *Pf*Atg8 and 70-74 in *Sc*Atg8, were also constructed. Strains expressing these constructs were designated *Pf*Atg8_*Sc*Loop and *Sc*Atg8_*Pf*Loop (Fig. 3C). Finally, we constructed hybrids of *Pf*Atg8 and *Cp*Atg8 whose loop regions, *Pf*Atg8 amino acids 69-82 and *Cp*Atg8 amino acids 69-86, were exchanged (*Pf*Atg8_*Cp*Loop and *Cp*Atg8_*Pf*Loop). All constructs were *N*-terminally tagged with GFP.

**Figure 3.**
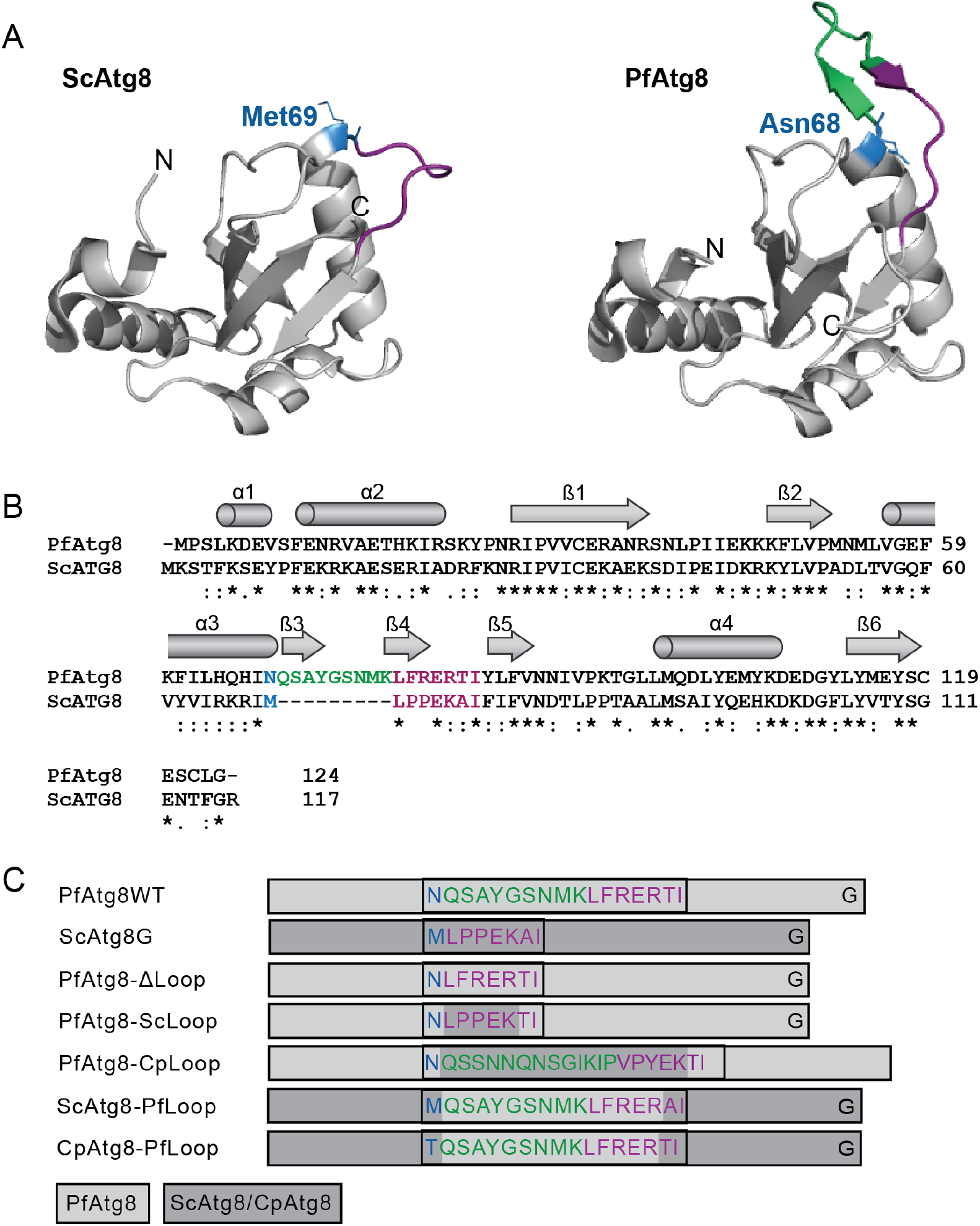
Constructs for testing the requirement for the Apicomplexan-specific loop in *Pf*Atg8. (A) Structures of *S. cerevisiae* Atg8 (3VXW) and *P. falciparum* Atg8 (4EOY). The region corresponding to the loop in canonical homologs is in purple. The additional 9 amino acid insertion in *Pf*Atg8 is in green. Amino acid residues preceding the loop region are shown as blue sticks. (B) Clustal Omega alignment of *S. cerevisiae* (YBL078C) and *P. falciparum* (PF3D7_1019900) Atg8 sequences. Color code the same as in (A). Secondary structure based on *Pf*Atg8 structure 4EOY are indicated. (C) Schematic representation of Atg8 hybrid constructs designed for testing structural requirements for Atg8 function in malaria parasites. Light grey rectangles denote *Plasmodium* sequence, dark grey rectangles denote *S. cerevisiae* or *C. parvum* sequence. The sequence of the loop region is colored as in (A). Note that all constructs were *N*-terminally tagged with GFP for easier detection which is not shown here for clarity.

When lysates from parasites expressing the hybrid constructs were subjected to Triton X-114 fractionation, *Pf*Atg8ΔLoop and the hybrids in which the entire *Plasmodium* loop region was replaced with the corresponding sequence from yeast or *Cryptosporidium* homologs (*Pf*Atg8_*Sc*Loop and *Pf*Atg8_*Cp*Loop) partitioned to the detergent fraction at 56%, 93% and 61% of endogenous Atg8 respectively, indicating that these hybrids are recognized by the *Plasmodium* conjugation machinery (Fig. 4A and B). Also *Cp*Atg8_*Pf*Loop, similar to the full length *Cp*Atg8, partitioned to the membrane fraction comparably to *Pf*Atg8WT. In contrast, *Sc*Atg8_*Pf*Loop failed to associate with the membranes. Triton X-114 fractionation results were corroborated by live fluorescence microscopy (Fig. 4C and D). GFP-tagged *Pf*Atg8ΔLoop, *Pf*Atg8_*Sc*Loop, *Pf*Atg8_*Cp*Loop, and *Cp*Atg8_*Pf*Loop but not *Sc*Atg8_*Pf*Loop localized to structures typical of the elongated, branched or segmented apicoplast with some cytosolic signal. *Sc*Atg8_*Pf*Loop showed cytosolic localization with occasional brighter patches that we cannot interpret. Taken together, in contrast to previously reported *in vitro* binding assays, we conclude that *in vivo*, the Apicomplexan-specific loop is not required for Atg8 recognition by the *Plasmodium* conjugation machinery.

**Figure 4.**
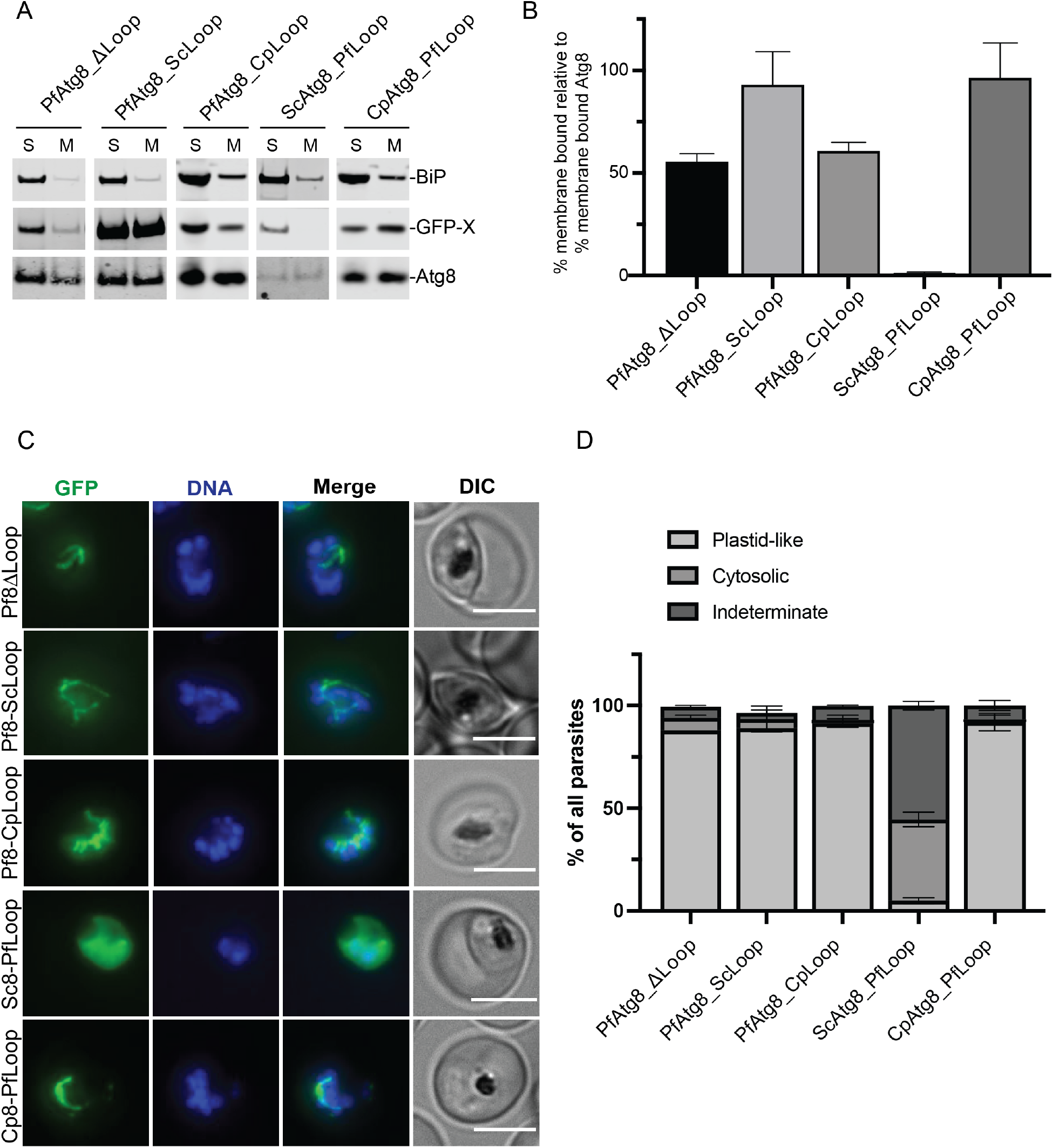
Apicomplexan-specific loop is not required for membrane conjugation of Atg8. (A) Western blot showing membrane association of indicated Atg8 mutants in a Triton X-114 fractionation experiment. S, soluble fraction; M, membrane fraction; BiP, soluble protein; Atg8, membrane-bound protein; X, Atg8 homolog. (B) Percentage of membrane-bound Atg8 mutants normalized to the percentage of the membrane-bound endogenous Atg8. Mean ± standard deviation from 2 replicates is shown. (C) Representative live microscopy images showing the localization of indicated Atg8 mutants. DNA was stained with Hoechst 33342. Maximum intensity projections are shown. Scalebar 5 μm. (D) Quantification of parasites with the indicated localization pattern of GFP-tagged Atg8 mutants. Mean ± standard deviation of 2 independent experiments is shown.

### Atg8 mutants are conjugated to the apicoplast membrane but do not have effector function

Most known Atg8 interactions including those with the membrane conjugation machinery and autophagic cargo adapters occur via the LIR-LDS interface (4). The LIR motif contains two hydrophobic amino acids separated by two random residues. These two hydrophobic residues bind to two hydrophobic pockets in the LDS on the Atg8 molecule (14, 30, 31). These interactions may be stabilized by additional contacts involving residues downstream or upstream of the LIR motif and residues outside the core LDS (3, 19, 32–34). A patch of positively-charged aminoacids, RKR or RRR, at the end of helix α3 in mammalian LC3 and GABARAP homologs has been implicated in binding to residues adjacent to LIR motifs and modulating selectivity and/or affinity of the LIR-LDS interactions (19, 35, 36). The corresponding region of *Pf*Atg8 (residues 66-68) is not positively-charged but instead consists of amino acids HQH (Fig. S1). We wondered whether this change was important for LIR-LDS interactions with the conjugation machinery or perhaps a new effector interaction. Therefore, we tested a *Pf*Atg8 mutant in which the residues HQH were replaced with RKR (*Pf*Atg8_RKR) for apicoplast membrane association and functional complementation. In the *Pf*Atg8 crystal structure, H66 of *Pf*Atg8 forms a hydrogen bond with the LIR motif of Atg3, in addition to the typical interaction within the hydrophobic pockets of the core LDS (29). Consistent with an interaction with membrane conjugation enzymes, *Pf*Atg8_RKR did not show membrane binding in a Triton-X114 fractionation assay, did not localize to the apicoplast and consistently failed to restore growth of parasites lacking Atg8 (Fig. 5A-D).

**Figure 5.**
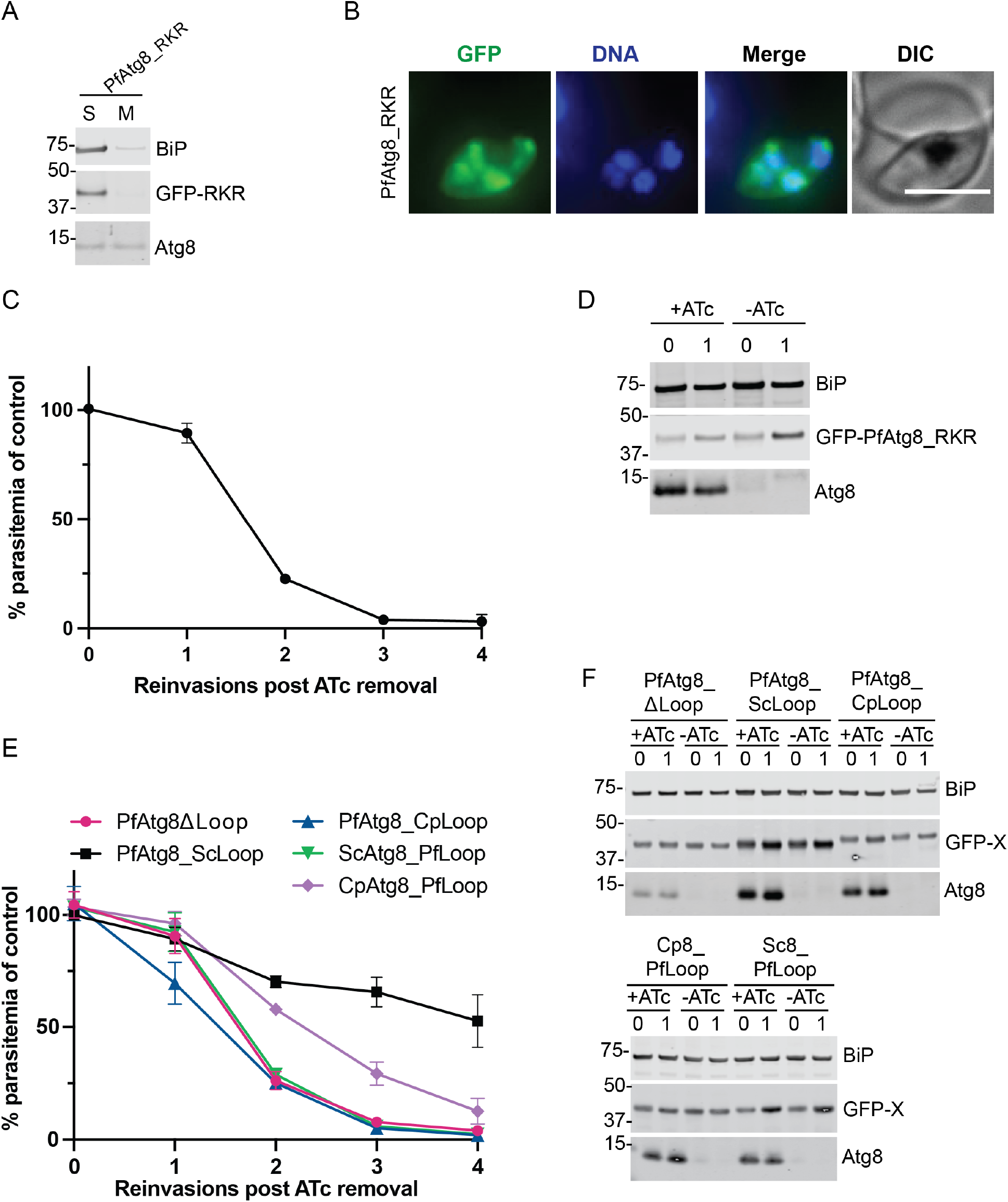
A region outside LDS and other than the Apicomplexan loop is required for *Pf*Atg8 effector function. (A) Western blot showing membrane association of *Pf*Atg8_RKR mutant in a Triton X-114 fractionation experiment. S, soluble fraction; M, membrane fraction; BiP, soluble protein; Atg8, membrane-bound protein; X, Atg8 homolog. (B) Representative live fluorescence images showing the localization of *Pf*Atg8_RKR mutant. DNA was stained with Hoechst 33342. Maximum intensity projections are shown. Scalebar 5 μm. (C) Growth of Atg8-TetR/DOZI parasites expressing GFP-*Pf*Atg8_RKR in the absence of ATc over 4 reinvasion cycles. Parasitemia for each time point was normalized to the culture grown in the presence of ATc. Mean ± standard deviation for 2 biological replicates is shown. (D) Western blots showing expression of the indicated proteins in the first two cycles of the growth time course. BiP, loading control; Atg8, endogenous Atg8. Numbers above the blot represent parasite reinvasion cycles post ATc removal. (E) Growth of Atg8-TetR/DOZI parasites expressing indicated GFP-tagged Atg8 mutants in the absence of ATc over 4 reinvasion cycles. Parasitemia for each time point was normalized to the culture grown in the presence of ATc. Mean ± standard deviation for 2 biological replicates is shown. (F) Western blots showing expression of the indicated proteins in the first two cycles of the growth time course. BiP, loading control; Atg8, endogenous Atg8; X, Atg8 mutant. Numbers above the blot represent parasite reinvasion cycles post ATc removal.

Notably, *Cp*Atg8 is conjugated to the apicoplast membrane (indicating an intact LDS-LIR interaction) but is unable to rescue growth defects in parasites lacking endogenous Atg8 (Fig. 1C-D and Fig. 2A-B). This result suggested that *Pf*Atg8 effector function requires regions outside the LDS. To date, very few effectors have been shown to interact with Atg8 proteins using alternative interaction sites (16, 17). We wondered if the Apicomplexan-specific loop might be involved in a new interaction for *Pf*Atg8 effector function. The Atg8 loop hybrids that were able to conjugate to the apicoplast membrane (see previous results section) were further tested for their ability to rescue growth defects in cells lacking endogenous Atg8. While *Pf*Atg8ΔLoop, *Pf*Atg8_*Cp*Loop and *Cp*Atg8_*Pf*Loop could not rescue growth inhibition in Atg8-deficient parasites, parasites expressing *Pf*Atg8_*Sc*Loop could replicate in the absence of endogenous Atg8, albeit at a slower rate than control parasites (Fig. 5D). Therefore, while *Pf*Atg8 effector function likely requires regions outside the LDS, the extended loop of *Pf*Atg8 was not strictly required. The Apicomplexan-specific loop may instead have a more general role, for example in maintaining protein stability.

### Role of previously-identified Atg8 effector regions in *Pf*Atg8 function

In addition to the LDS, other regions of Atg8 have been implicated in intermolecular interactions of canonical Atg8 homologs. The *N*-terminal helical region, which generally has little sequence conservation between homologs, has been implicated in oligomerization, binding of tubulin and lipids, and membrane tethering and fusion. It is also thought to contribute to selectivity of LIR-LDS interactions (2, 15, 16, 37). To test whether this region is required for *Pf*Atg8 function, we replaced the *N*-terminal helix *α*1 of *Pf*Atg8 (amino acids 1-9) with the corresponding region from yeast Atg8 (amino acids 1-10) and transfected it into Atg8-TetR/DOZI parasites. This construct, named *Sc*Helix1, correctly localized to the apicoplast, was membrane bound and rescued parasite growth upon depletion of endogenous Atg8 (Fig. 6A-C). We also attempted to replace both *N*-terminal helices with the corresponding region of yeast Atg8 but never obtained transfectants.

**Figure 6.**
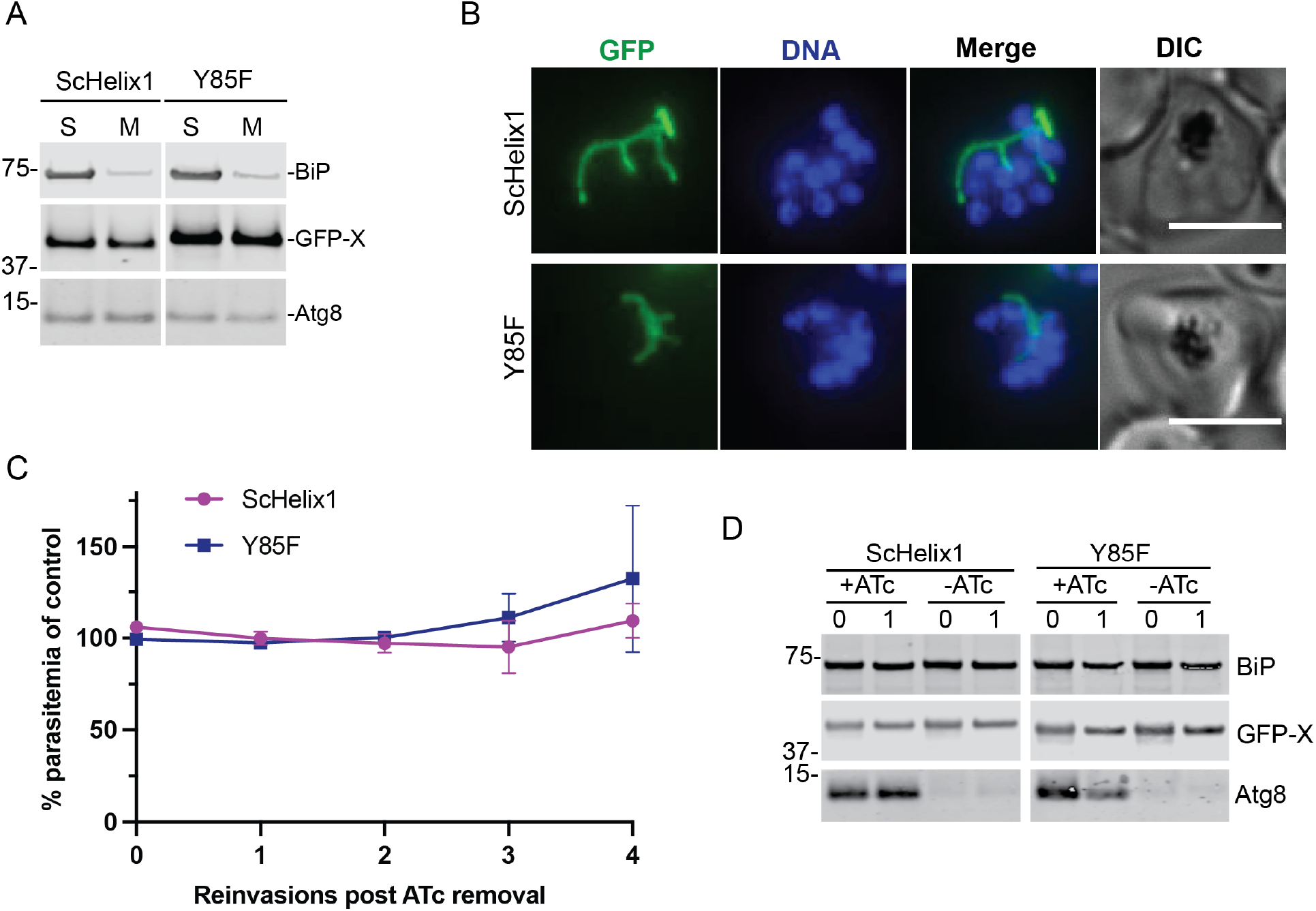
Previously identified interaction sites on Atg8 are not required for *Pf*Atg8 apicoplast function. (A) Western blot showing membrane association of *indicated Pf*Atg8 mutant in a Tritont X-114 fractionation experiment. S, soluble fraction; M, membrane fraction; BiP, soluble protein; Atg8, membrane-bound protein; X, Atg8 homolog. (B) Representative live fluorescence images showing the localization of the indicated *Pf*Atg8 mutants. DNA was stained with Hoechst 33342. Maximum intensity projections are shown. Scalebar 5 μm. (C) Growth of Atg8-TetR/DOZI parasites expressing indicated GFP-tagged *Pf*Atg8 mutants in the absence of ATc over 4 reinvasion cycles. Parasitemia for each time point was normalized to the culture grown in the presence of ATc. Mean ± standard deviation for 2 biological replicates is shown. (D) Western blots showing expression of the indicated proteins in the first two cycles of the growth time course. BiP, loading control; Atg8, endogenous Atg8. Numbers above the blot represent parasite reinvasion cycles post ATc removal.

A recently identified <u>U</u>biquitin interacting motif (UIM) <u>D</u>ocking <u>S</u>ite (UDS) serves as a binding site for proteins containing UIM motif, including a number of autophagy receptors and the Atg4 protease (17, 38, 39). Its consensus sequence is Ψ-F-Ψ-Ω/T, where Ψ is hydrophobic and Ω is aromatic. The second residue, Phe, is highly conserved in most canonical homologs and is critical for interactions with the effectors and cannot be mutated. *Pf*Atg8, as well as its homologs or putative homologs from other organisms containing a secondary plastid, has a tyrosine residue in this position (Tyr85). We therefore replaced Tyr85 with Phe and episomally expressed it as a GFP fusion in Atg8-TetR/DOZI parasites. This mutant, denoted as Y85F, also bound membranes, showed apicoplast localization and restored growth of parasites lacking endogenous Atg8 (Fig. 6A-C).

In conclusion, neither the most *N*-terminal helix nor the UDS region are specifically required for Atg8 function in *Plasmodium* apicoplast.

### New tools for studying apicoplast biogenesis

One of the challenges we encountered during our work on *Pf*Atg8 function was the reliable expression of tagged Atg8 constructs with the correct expression pattern. Atg8 mRNA levels are low until they peak in the late stages of *Plasmodium* cell cycle (40-48 hrs), an expression pattern that cannot be recapitulated with commonly used promoters (40). Expression from episomes also results in highly variable protein levels with heterogeneity in the cell population. Finally, the *C*-terminus of Atg8 must be unmodified because it is required for the covalent attachment to the membrane which adds difficulties to modifications at the endogenous locus (41). Correct expression timing or protein levels may affect interpretation of the data; in fact, our episomally expressed GFP-Atg8 fusions often had low expression levels in schizont parasites, which made it difficult to see the localization on segmented apicoplasts, whereas in earlier trophozoite stages we frequently saw a large GFP-labeled punctate structure with additional cytosolic signal.

To facilitate studies of *Pf*Atg8 and the apicoplast biogenesis we generated two *Plasmodium* strains in which Atg8 was *N*-terminally tagged at its endogenous locus. The first strain contains GFP-Atg8 in Atg7-TetR/DOZI background (Fig. 7A). For its generation we utilized CRISPR/Cas9 system to introduce the *N*-terminal GFP tag on Atg8 (42), and fluorescence activated cell sorting (FACS) to select parasites in which GFP was integrated. By using FACS, we bypassed the need for using drug selection; this leaves one more selection marker for introducing additional constructs. Downregulation of Atg7 expression blocks membrane conjugation of Atg8 leading to defects in apicoplast biogenesis and parasite growth inhibition, which is rescued by addition of IPP (26, 41) (Fig. S3).

**Figure 7.**
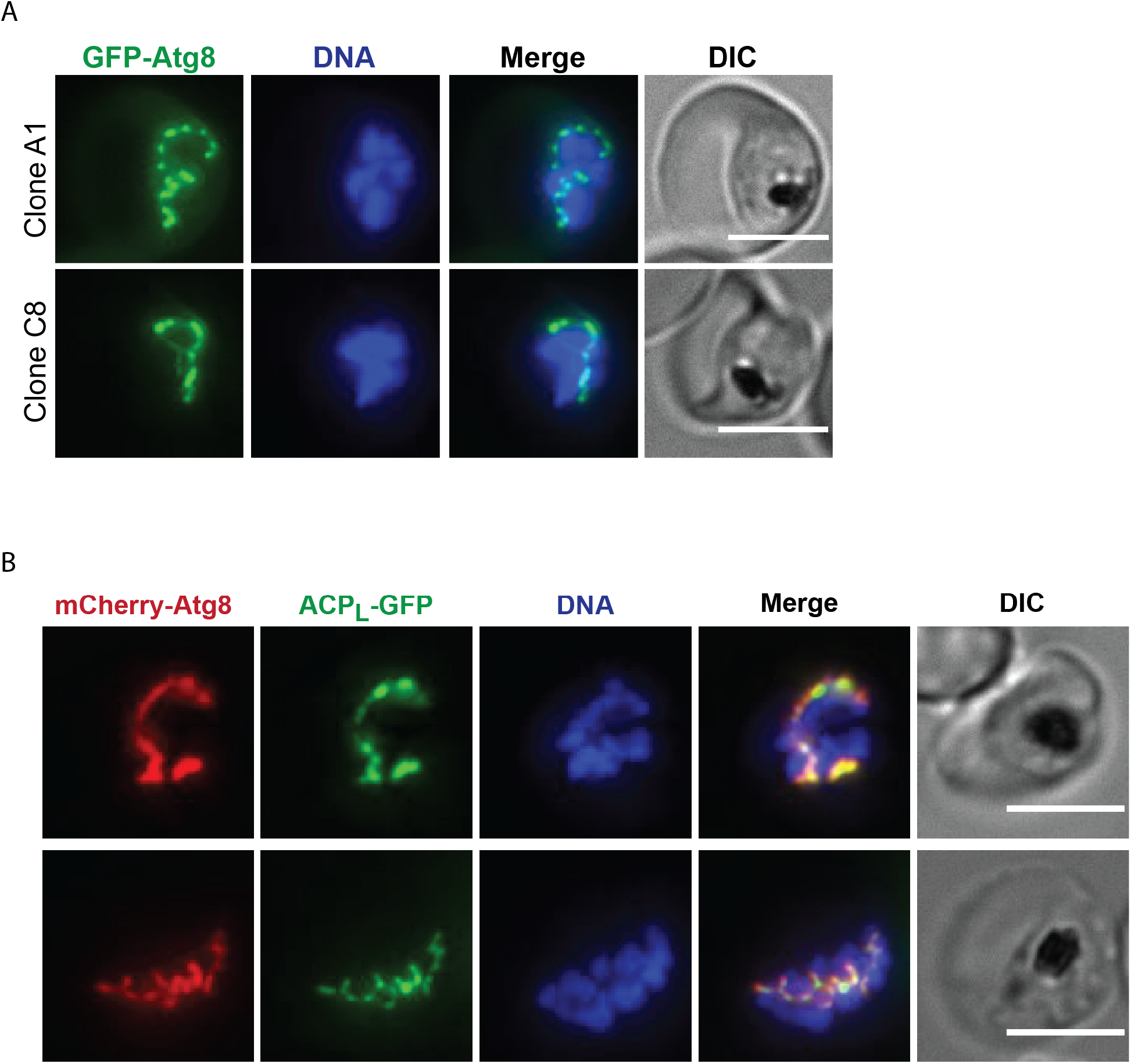
Strains expressing Atg8 fluorescently tagged at the endogenous locus. (A) GFP-Atg8 expression in Atg7tetR/DOZI parasites. Representative live fluorescence images of two clonal lines are shown. (B) Representative live fluorescence images of parasites expressing ATc-regulatable mCherry-Atg8 and episomal ACP_L_-GFP. Maximum intensity projections are shown. Scalebar 5 μm.

The second strain contains two apicoplast markers: (1) Atg8, which labels outer apicoplast membrane and was modified with an *N*-terminal mCherry tag and an aptamer array in the 3’ untranslated region for ATc-mediated regulation of expression (24) (Fig. S3); and (2) episomally expressed GFP targeted to the apicoplast via the ACP leader sequence, ACP_L_-GFP, which marks the apicoplast lumen(13) (Fig. 7B). Having these two markers expressed in one cell could be particularly useful for tracking the behavior of both the membrane and the lumen of the apicoplast during the division and segregation. We have occasionally seen that GFP-Atg8 localized to structures resembling a necklace of beads pattern or a comb-like structure (see Fig. 7A and Fig. 4C, *Pf*_*Cp*Loop). This may be an intermediate product of apicoplast division worth studying more in detail using high resolution microscopy techniques.

Overall, these two strains offer the possibility to study apicoplast biogenesis using Atg8 expressed at endogenous levels at the right stage of the parasite’s cell cycle, bypassing the need for generating anti-Atg8 antibodies.

## Discussion

Atg8 family proteins are critical players in the cell’s endomembrane system with diverse functions in autophagy and non-autophagic pathways (3, 43). However, little is known about the molecular basis for Atg8 functional diversification, particularly for non-autophagic functions. In *Plasmodium*, the single Atg8 ortholog has evolved an essential function in the biogenesis of the apicoplast, a unique plastid of secondary endosymbiosis origin (10, 13). Here we used a functional complementation approach to determine structural requirements for this unusual function of *Pf*Atg8. We showed that the new effector function of *Pf*Atg8 requires interactions other than those previously identified for macroautophagy and selective autophagy functions: An intact LDS sufficient for interaction with membrane conjugation machinery was nonetheless insufficient to restore apicoplast effector function. Mutation of another previously-identified interaction surface UDS or the *N*-terminal helix also did not affect apicoplast effector function. Finally, even the Apicomplexan-specific loop was not required for membrane binding nor effector function *in vivo*.

Atg8 proteins and their conjugation machinery are highly conserved among eukaryotes, and it is presumed that a pathway involving Atg8 was present in the “last eukaryotic common ancestor,” or LECA (5). The secondary endosymbiosis origin of the apicoplast indicates that the apicoplast function was a later adaptation of Atg8 proteins. However, an intriguing question remains: was autophagy or another membrane-related pathway the ancestral function of Atg8? The pathways in which Atg8 family proteins function ultimately involve transport of material in a membrane vesicle between membrane compartments in the endomembrane system, including the ER, Golgi, plasma membrane and the vacuole/lysosome (44–51). However, the functional diversification of Atg8 proteins beyond autophagy is most apparent in organisms that possess multiple Atg8 orthologs like mammals. Instead, most eukaryotes from fungi, protists, sponges to higher multicellular organisms, encode at least a minimal set of Atg proteins and have a version of autophagy or autophagy-like pathway manifested by the formation of double membrane vesicles (12, 52–56). Based on this evolutionary pattern, autophagy was most likely the ancestral function of Atg8 and subsequent environmental adaptations led to diversification of Atg8 function through expansion, reduction, or complete loss of autophagy machinery (12, 57). Studies of Atg8 in a greater diversity of organisms will provide more support for the evolutionarily ancestral function of Atg8.

If we assume autophagy was the ancestral function of Atg8 proteins, it is intriguing how such a compact protein (<150 amino acids) proceeded to evolve so many diverse non-autophagic functions. Atg8 proteins generally function as membrane-bound scaffolds that recruit other factors. We hypothesize that *Pf*Atg8 acts in a similar manner, i.e. by recruiting other proteins that in turn carry out the effector functions. Most known Atg8 interactions occur via LDS on Atg8 which bind LIR motifs of the other proteins (4). Additionally, the N-terminal helical region which distinguishes Atg8 proteins from other ubiquitin-like proteins, and UDS which is located on the side of Atg8 molecule opposite to LDS have also been implicated in Atg8 interactions (16, 17). Our results indicate that none of these three regions is sufficient to confer *Pf*Atg8’s apicoplast function, suggesting that another distinct site is involved.

The most obvious candidate was the loop between the helix α3 and the β-strand β5 in *Pf*Atg8 structure. This loop is longer in *Pf*Atg8 and its homologs from organisms harboring a secondary plastid than in canonical Atg8 proteins. Surprisingly, this extended loop is also not strictly required for the effector function because when replaced with the corresponding (shorter) sequence from yeast, such Atg8 mutant can still partly rescue parasites lacking endogenous Atg8. Its replacement with *Cryptosporidium* loop or deletion of the additional 9 amino acids in the *Plasmodium* loop renders the protein non-functional. Is it required for another function like stabilizing the protein structure? And most importantly, what are the structural features responsible for the effector function of *Pf*Atg8? Are they located in the *C*-terminal part downstream of UDS that has not been investigated?

Structural data are needed to answer these questions and better understand the structure-function relationship for the unique role of Atg8 in *Plasmodium*. We attempted to crystallize *Toxoplasma* and *Cryptosporidium* Atg8 homologs as well as the loop mutants of *Pf*Atg8, unfortunately our attempts were so far unsuccessful. Identifying effectors that interact with *Pf*Atg8 on the apicoplast will also provide insights into the molecular mechanism of Atg8 in apicoplast biogenesis. Strains generated in this study expressing Atg8 mutants and homologs that correctly localize to apicoplast membrane but lost the effector function could facilitate identification of Atg8 effectors by co-IP/MS. Finally, *Plasmodium* strains with endogenously fluorescently tagged Atg8 will help visualization of apicoplast biogenesis by microscopy.

## Materials and methods

### Cloning

Primers and gBlocks used in this study are listed in Supplementary Table S1. All constructs for genetic complementation were cloned into pfYC110 using In-Fusion. Atg8 homologs were cloned without the C-terminal extension following the last glycine residue. To clone pfYC110-GFP-*Pf*Atg8WT, GFP with a Gly-Ala-Gly-Ala linker and AatII site was amplified from pfYC110-ACP_L_-GFP and *Pf*Atg8 was PCR-amplified from *Plasmodium* genomic DNA. The two fragments were inserted into pfYC110, resulting in plasmid pfYC110-GFP-*Pf*Atg8WT. *Pf*Atg8 was removed from that plasmid by AatII-SacII digest and replaced with Atg8 homologs. Human Atg8 homologs LC3A, LC3B, GABARAP and GABARAPL1 were amplified from cDNA libraries kindly provided by S. Pfeffer and L. Li labs; gBlocks (IDT DNA) were used for cloning human LC3C and GABARAPL2; *Sc*Atg8G was amplified from yeast S288C genomic DNA. To clone the *Plasmodium*-yeast loop hybrids, parts of Atg8 from the start codon to the loop or from the loop to the stop codon were amplified from the plasmids carrying *Pf*Atg8 or *Sc*Atg8 and the desired loop sequence was added on the primer; the *N-* and *C-*terminal parts of the hybrid were then fused by overlap extension PCR; the N-terminal GFP tag was amplified with a Gly-Ala-Gly-Ala linker and PvuI site downstream of it; both fragments were simultaneously inserted into pfYC110 via AvrII-SacII restriction sites. The remaining hybrid constructs as well as CpAtg8 were purchased as gBlocks (IDT DNA) and inserted into pfYC110 via AvrII-SacII sites.

Accession numbers: hLC3A isoform 1, NP_115903; hLC3B, NP_073729; hLC3C, NP_001004343; hGABARAP, NP_009209; hGABARAPL1 isoform 1, NP_113600; hGABARAPL2, NP_009216; *Pf*Atg8, PF3D7_1019900; *Tg*Atg8, TGME49_254120; *Cp*Atg8, CPATCC_0009830.

To clone the construct for the endogenous tagging of Atg8 (pSN_054_GFP-Atg8-gRNA), the left and right homology regions were amplified from *Plasmodium* genomic DNA; GFP followed by a Gly-Ala-Gly-Ala linker was amplified from pfYC110-GFP-*Pf*Atg8WT. Left homology region contains the sequence directly preceding Atg8 ORF whereas the right homology region consists of Atg8 ORF and 423 bp downstream of it. Primers introduced restriction sites XmaI and ApaI right upstream and downstream of GFP and silent mutations in Atg8 ORF in the region corresponding to gRNA. The three products were inserted into the linear plasmid pSN_054_V5 (a gift from J. Niles) via FseI and SacII using Gibson assembly. Next, gBlock15 containing the guide RNA expression cassette for CRISPR/Cas9 editing was inserted into the SacII site. The gRNA was designed using http://grna.ctegd.uga.edu/.

To generate the plasmid pSN_054_mCherry-Atg8-TetR/DOZI, left homology region was amplified from pSN_054_GFP-Atg8-gRNA; mCherry was amplified from pLN-mCherry-T2A-ACP_L_-GFP (26). Atg8 ORF containing silent mutations in the region complementary to gRNA and an N-terminal Gly-Ala-Gly-Ala linker was amplified from pSN_054_GFP-Atg8-gRNA. The mCherry tag and Atg8 ORF were then stitched together by overlap extension PCR. The resulting product and the left homology region were inserted into pSN_054 linearized with FseI and ApaI, resulting in an intermediate plasmid pSN_054_LeftSide. Next, the gRNA expression cassette was amplified from pSN_054_GFP-Atg8-gRNA and the right homology region was amplified from pSN053-ATG8 (13). Both products were inserted into pSN_054-LeftSide via AscI and I-SceI restriction sites.

### Culture and transfection

*Plasmodium falciparum* parasites were grown in human erythrocytes (Stanford Blood Center, Stanford, CA) at 2% hematocrit under 5% O_2_ and 5% CO_2_, at 37°C in RPMI 1640 medium supplemented with 5 g/L AlbuMAX II (Gibco), 2 g/L NaHCO_3_ (Fisher), 25 mM HEPES (pH 7.4) (Sigma), 0.1 mM hypoxanthine (Sigma), and 50 mg/liter gentamicin (Gold Biotechnology) (further referred to as culture medium). For transfections, 200 µl packed red blood cell per transfection were washed twice in cytomix, combined with 50-100 µg ethanol-precipitated plasmid DNA dissolved in 30 μl TE buffer and to 170 µl cytomix and tranferred to a 0.2 cm electroporation cuvetter. Erythrocytes were electroporated at 310V, 950 µF, infinite resistance using a Gene Pulser Xcell electroporation system (BioRad). Electroporated red blood cells were added to schizont stage parasites from 0.5 ml culture at 5% parasitemia, 2% hematocrit. For drug selection, the following drugs were used as applicable starting 3 days after transfection: 2.5 mg/L blasticidin S (RPI Research Products), 2.5 nM WR99210, 500 µg/ml G418 sulfate (Corning); 0.5 µM anhydrotetracycline (Sigma) was added to parasites to maintain expression of TetR/DOZI regulated genes.

For selection of endogenously tagged parasites which did not have a drug selection marker, transfections were performed as described above and a pool of GFP-positive transfected parasites was isolated 4 days post transfection using Sony SH800S Cell Sorter. Parasites were fed with culture medium supplemented with 0.5 µM ATC for two weeks followed by another FACS sort to isolate single clones. To set gates for FACS sorting, the following cells were used: uninfected red blood cells and red blood cells infected with non-fluorescent parental Atg7-TetR/DOZI strain (background fluorescence), red blood cells infected with Atg7-TetR/DOZI episomally expressing GFP-Atg8 (GFP fluorescence).

### Growth assays

Ring-stage parasites at 5 to 10% parasitemia were washed twice in the culture medium to remove ATc and resuspended in the culture medium, and the hematocrit was adjusted to 2%. Parasites were divided into 2 cultures grown in the culture medium supplemented with 0.5 µM ATc, or without ATc. For validating strains with endogenously tagged Atg8, a third condition without aTC with 200 µM IPP (Isoprenoids LLC) was included. The growth assays were continued for 4 reinvasion cycles. At the schizont stage of each cycle, parasitemia was measured by flow cytometry (BD Accuri), starting at 24 hrs (0 reinvasions) post ATc removal. Culture aliquots were incubated with 17 μM dihydroethidium (Thermo Fisher) for 15 min at RT to stain nuclei and 50,000 events were analyzed on a BD Accuri C6 flow cytometer. Cultures were diluted with fresh medium with red blood cells at 2% hematocrit so that the parasitemia in +ATc condition is 1% and the -ATc condition was diluted by the same factor; ATc or IPP was added as required. Aliquots of culture for western blotting were taken at 24 hrs and 72 hrs (0 and 1 reinvasion) post ATc removal.

### Western blot

Erythrocytes were lysed with 0.1% saponin for 5 min on ice to release parasites. Parasite pellets were washed twice with ice-cold PBS, resuspended in 2× lithium dodecyl sulfate (LDS) buffer (Life Technologies) in PBS supplemented with 25 µM DTT and boiled 5 min at 96 °C. After separation on Bis-Tris Novex gels (Invitrogen), proteins were transferred to a nitrocellulose membrane, blocked with a blocking buffer containing 0.1% Hammarsten grade casein (Affymetrix), 0.2× PBS, and 0.01% sodium azide. Next, membranes were incubated with primary antibodies diluted in a wash buffer consisting of 50% blocking buffer, 50% Tris-buffered saline and 0.25% Tween (TBST) overnight at 4°C or 1 h at room temperature, followed by 3 washes with TBST and 1 h incubation with the secondary antibodies diluted in the same buffer. Blots were visualized using the LiCor double-color detection system and converted to grayscale images for the purpose of this publication. Primary antibodies and dilutions used in this study: guinea pig anti-*Pf*Atg8 serum (Josman LLC), 1:1,000; mouse anti-GFP (Clontech, 632381), 1:10,000; mouse anti-*Plasmodium yoelli* BiP (a gift from Sebastian Mikolajczak and Stefan Kappe), 1:20,000. Secondary fluorophore-conjugated antibodies were purchased from Fisher (LiCor) and used at 1:10,000 dilution.

### Triton X-114 fractionation assay

Schizont stage parasites were lysed with 0.1% saponin and washed 3 times with ice-cold PBS. Parasite pellets were resuspended in ice-cold lysis buffer (1× PBS, 1% Triton X-114 [Thermo Scientific 28332], 2 mM EDTA, 1× protease inhibitors [Pierce A32955]) and incubated on ice for 30 minutes. Cell debris were removed by 10-minute centrifugation at 16,000 × g, 4°C. Supernatant was transferred to a fresh Eppendorf tube, incubated 2 minutes at 37°C to allow phase separation, and centrifuged 5 minutes at 16,000 × g at room temperature. The top (aqueous) layer was transferred to another tube. The interphase was removed to avoid cross-contamination between the layers. The bottom (detergent) layer was resuspended in 1× PBS, 0.2 mM EDTA to equalize the volumes of the two fractions. Both fractions were subjected to methanol-chloroform precipitation, resuspended in PBS containing 2× NuPAGE LDS sample buffer, boiled for 5 minutes at 95°C, and analyzed by western blot. Quantification of band intensities was done using ImageStudioLite. The percentage of each GFP fusion protein that was membrane bound was calculated and normalized to the percentage of endogenous Atg8 in the membrane.

### Microscopy

Live parasites in PBS with 0.4% glucose were incubated with 2 µg/ml Hoechst 33342 stain (Thermo Fisher H3570) for 15 min at room temperature to visualize nuclei. Images were acquired using the Olympus IX70 microscope equipped with a DeltaVision Core system, a 100× 1.4 NA Olympus lens, a Sedat Quad filter set (Semrock), and a CoolSnap HQ charge-coupled device (CCD) camera (Photometrics) controlled via SoftWoRx 4.1.0 software. Images were acquired as Z-stacks and analyzed using ImageJ. Unless stated otherwise, maximum intensity projections are shown in the figures.

## Acknowledgements

We thank Lingyin Li and Suzanne Pfeffer for human cDNA libraries, Sebastian Mikolajczak and Stefan Kappe for the anti-PyBiP antibody, and Aaron Straight for sharing the equipment.

This work was supported by the following grants to Ellen Yeh: NIH 1R01AI141366, Burroughs Wellcome Fund, Chan Zuckerberg Biohub. The funders had no role in study design, data collection and interpretation, or the decision to submit the work for publication.

## Supplemental material

**Figure S1.**
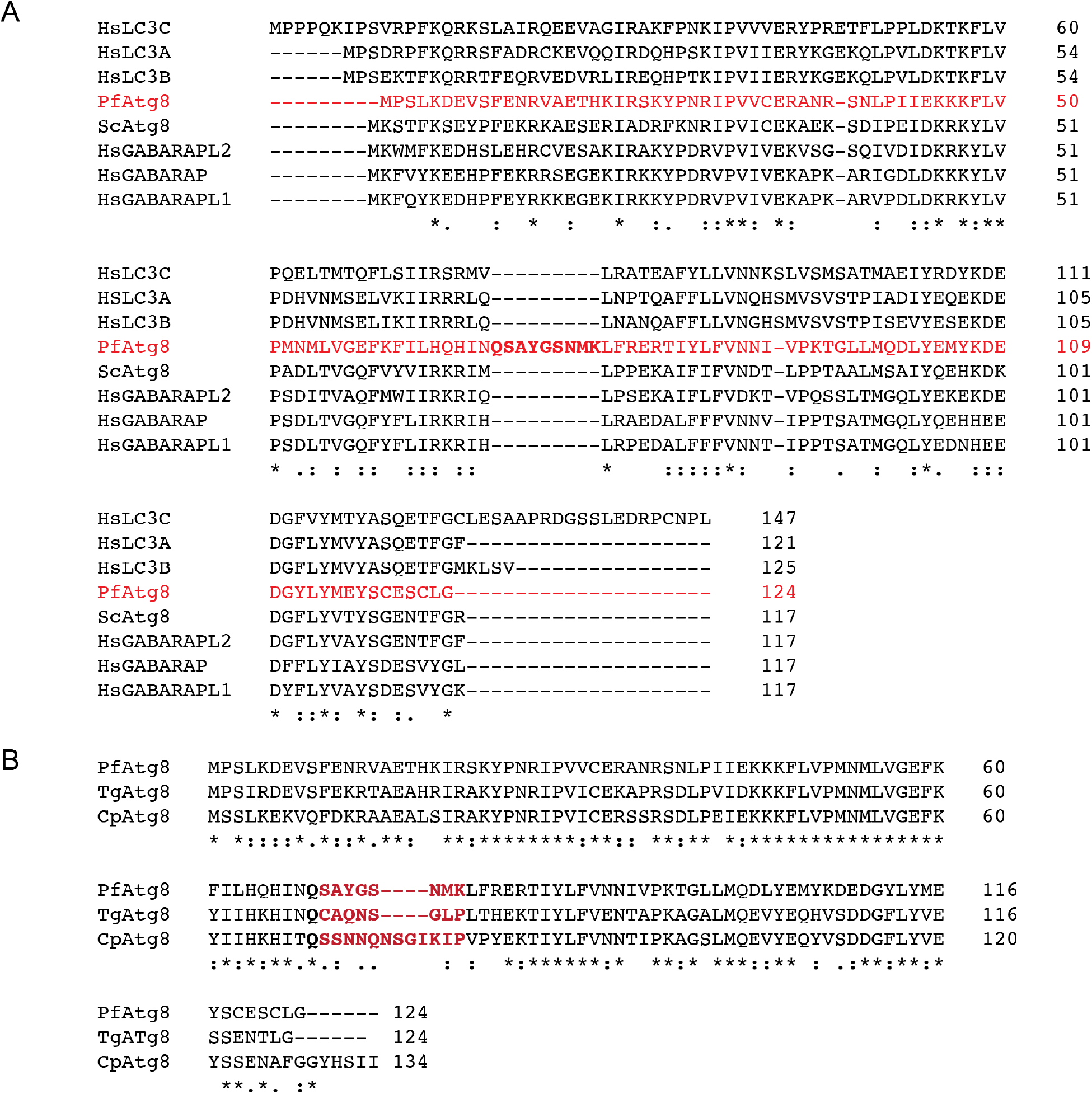
(A) Clustal alignment of *P. falciparum* (*Pf*), human (*Hs*) and yeast (*Sc*) Atg8 homologs. The additional amino acids in the Apicomplexan-specific loop of *Pf*Atg8 are highlighted in bold. (B) Clustal alignment of *P. falciparum* (*Pf*), *T. gondii* (*Tg*) and *C. parvum* (*Cp*) Atg8 homologs. The Apicomplexan-specific loop is highlighted in bold red. For accession numbers of the aligned sequences see *Cloning* section in Materials and Methods.

**Figure S2.**
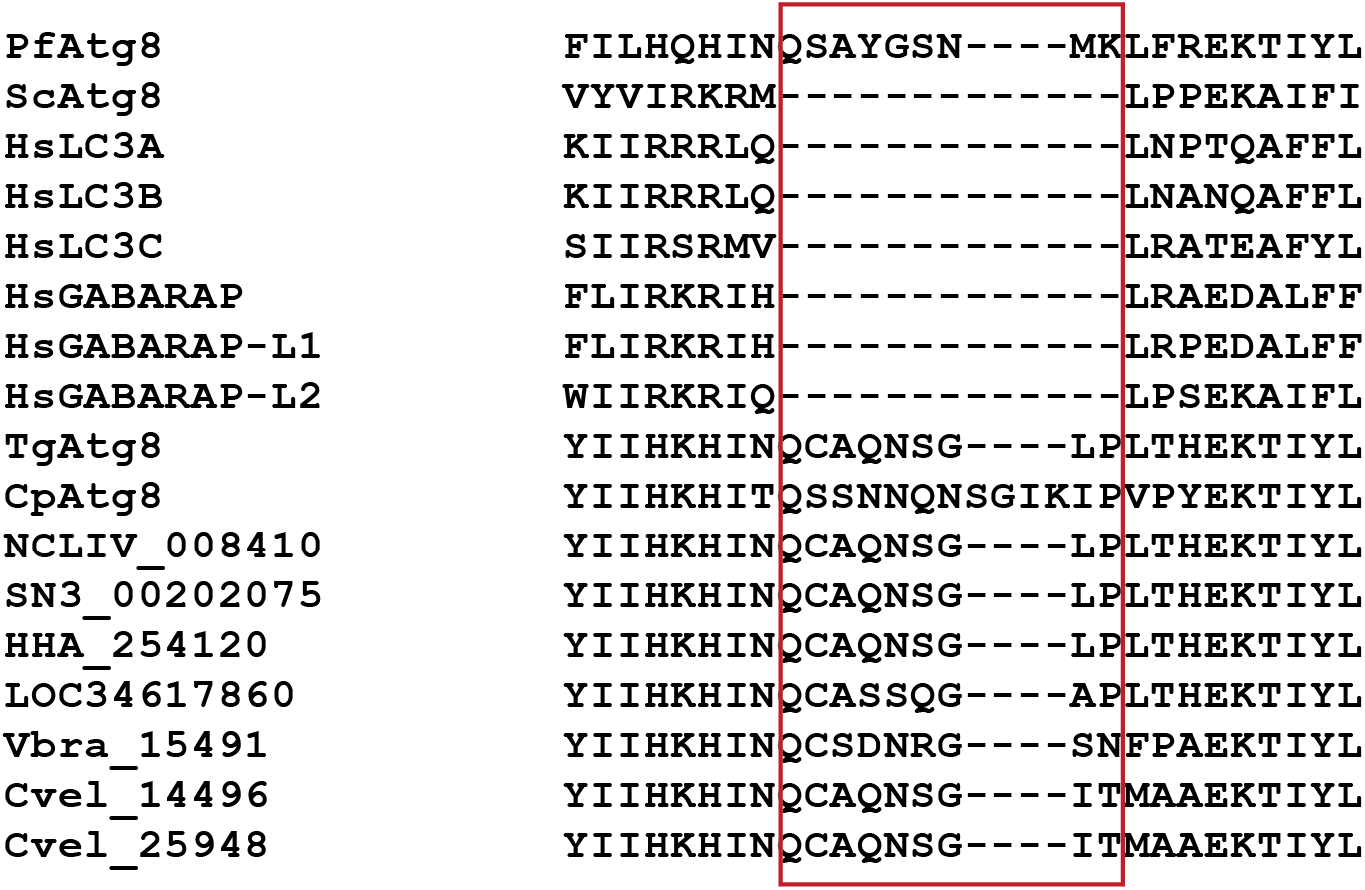
Clustal alignment of the loop region and neighboring amino acids in Atg8 homologs and putative homologs from *P. falciparum* (*Pf*), human (*Hs*), yeast (*Sc*), *T. gondii* (*Tg*), *C. parvum* (*Cp*), other apicomplexan parasites (*N. caninum*, NCLIV_008410; *S. neurona*, SN3_00202075; *H. hammondii*, HHA_254120; *C. cayetanensis*, LOC34617860), and chromerids containing a secondary plastid (*V. brassicaformis*, Vbra_15491; *C. velia*, Cvel_14496 and Cvel_25948). The additional amino acids absent from canonical homologs are in a red box.

**Figure S3.**
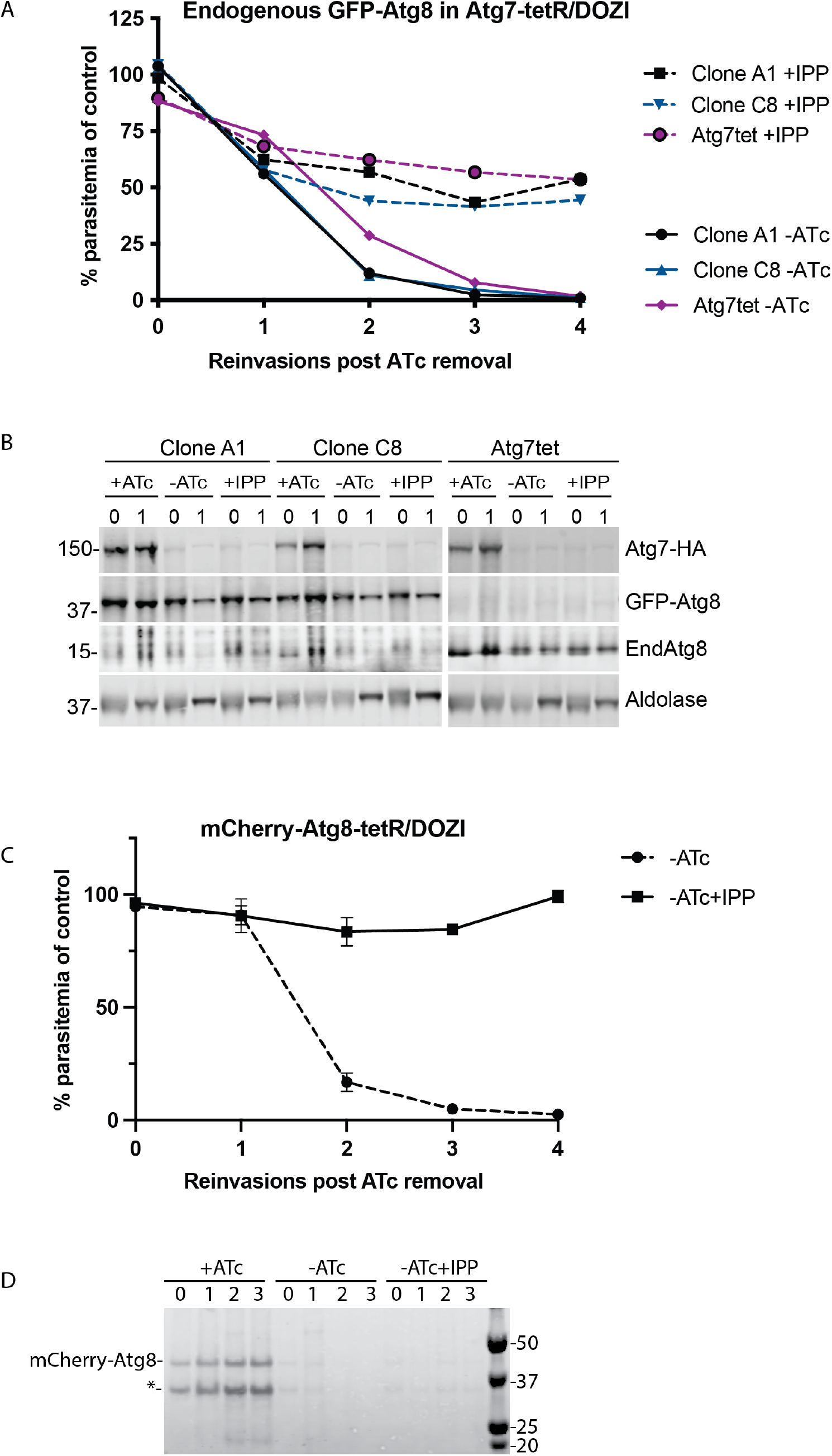
(A) Growth of indicated Atg7tetR/DOZI strains expressing endogenously tagged GFP-Atg8 and the parental Atg7-tetR/DOZI strain in the absence of Atg7(-ATc), with or without IPP over 4 reinvasion cycles. Parasitemia for each time point was normalized to the culture grown in the presence of ATc. (B) Western blot showing the expression of indicated proteins in the first two reinvasion cycles of the growth assay from (A). (C) Growth of mCherry-Atg8-tetR/DOZI in the absence of ATc, with or without IPP over 4 reinvasion cycles. Parasitemia for each time point was normalized to the culture grown in the presence of ATc. Mean ± standard deviation for 2 biological eplicates is shown. (D) Western blot showing the expression of mCherry-Atg8 in 4 reinvasion cycles of a growth assay shown in (C). Asterisk, unspecific band or degradation product of mCherry-Atg8

Table S1. Primers and gBlocks used in this study.

## Notes

### Competing Interest Statement

The authors have declared no competing interest.

